# Learning to Fold RNAs in Linear Time

**DOI:** 10.1101/852871

**Authors:** F A Rezaur Rahman Chowdhury, He Zhang, Liang Huang

## Abstract

RNA secondary structure is helpful for understanding RNA’s functionality, thus accurate prediction systems are desired. Both thermodynamics-based models and machine learning-based models have been used in different prediction systems to solve this problem. Compared to thermodynamics-based models, machine learning-based models can address the inaccurate measurement of thermodynamic parameters due to experimental limitation. However, the existing methods for training machine learning-based models are still expensive because of their cubic-time inference cost. To overcome this, we present a linear-time machine learning-based folding system, using recently proposed approximate folding tool LinearFold as inference engine, and structured SVM (sSVM) as training algorithm. Furthermore, to remedy non-convergence of naive sSVM with inexact search inference, we introduce a max violation update strategy. The training speed of our system is 41× faster than CONTRAfold on a diverse dataset for one epoch, and 14× faster than MXfold on a dataset with longer sequences. With the learned parameters, our system improves the accuracy of LinearFold, and is also the most accurate system among selected folding tools, including CONTRAfold, Vienna RNAfold and MXfold.

## 1 Introduction

For past decades, our understanding of ribonucleic acid (RNA) is changing. Proofs reveal that RNAs are involved in multiple processes, including gene expression, RNA modifications guiding [8] and particular diseases regulating [13]. RNA’s functionalities are highly related to its secondary structures, but determining the secondary structure using experimental methods is expensive, time-comsuming and difficult. Therefore, being able to rapidly and accurately predict RNA secondary structures is very useful and desired.

Accurate prediction model, i.e., well-designed features (for example base pair CG and AU, terminal mismatches, etc) and their weights, is one of the keys for solving RNA secondary structure prediction problem. Both thermodynamics-based models and machine learning-based models have been proposed and used for this problem, they share similar features, but use different ways to get feature weights. Thermodynamics-based models get feature weights directly from experimentally estimated thermodynamic parameters, and these models are widely used by RNA secondary structure prediction engines, such as Vienna RNAfold [15] and RNAstructure [17]. Thermodynamics-based models and engines constitute the most popular way for secondary structure prediction, however, they may suffer from the inaccurate measurement of thermodynamic parameters due to experimental limitation, which leads to an inaccurate prediction result [6].

Alternatively, machine learning-based models borrow feature templates from thermodynamics-based models, but use machine learning techniques to learn feature weights from known structures. The first trials are to borrow stochastic context-free grammars (SCFGs) learning framework [7,11,12] from natural languaging processing field, but due to the weak feature expression ability of SCFGs, their accuracies are all lower than thermodynamics-based models. To further improve accuracy, CONTRAfold [6] proposes a new machine learning-based method using Conditional Random Field (CRF) [14] for training, and achieves higher accuracy than thermodynamic methods. However, CONTRAfold training is very slow due to CRF algorithm, which makes it impossible for training on big dataset. A more recent machine learning-based work, MX-fold [1], uses structured SVM [19] instead of CRF to accelarate the training process. Though it is faster than CONTRAfold training, it borrows the inference implementation from CONTRAfold, which runs in cubic time and results in a costly inference process, especially when the training set includes long sequences. In addition, MXfold is not a pure machine learning-based model since it integrates thermodynamic model with learned model.

To overcome the efficiency bottleneck of training, we present a new machine learning-based RNA folding system, using the recently proposed linear-time prediction engine LinearFold [10] for inference. We use structured SVM algorithm for training. However, using LinearFold as inference may result in invalid updates due to the inexact search nature of LinearFold, and break the convergence property in naive sSVM training process. To remedy this, we utilize a max violation update strategy originally from violation-fixing structured perceptron [3,4,9], and generalize the naive sSVM algorithm to an advanced version of sSVM.

The results show that the training speed of our system is 41× faster than CONTRAfold for one epoch on a diverse dataset (average length 208.6 *nt*). On a dataset with more long sequences, our system is 14× faster than MXfold (average length 2713.5 *nt*). With the learned parameters, our system is the most accurate system among selected folding tools, including CONTRAfold, Vienna RNAfold and MXfold. Using a cross validation training, Our system improves the off-the-shelf LinearFold-C by +4.62% in PPV and +8.55% in sensitivity. Compared with retrained MXfold, our system gains +3.51% and +4.70% on PPV and sensitivity, separately.

Table 1 summarize the differences between 3 systems. Our contributions are as followed:

– We propose a fast training system which can do training and inference both in linear time. Our system can make big dataset training and long sequences training much more efficient.
– With the learned parameters, our system is more accurate compared with existing systems.
– We prove that sSVM converges when doing exact search for inference, but the convergence property does not hold when the search is inexact. We use a max violation update strategy to solve the convergence problem.

**Table 1:** Comparison of three machine learning-based RNA folding systems.

| *System* | Training |  |  |
| --- | --- | --- | --- |
|  | *Method* | *Inference Algorithm* | *Speed* |
| CONTRAFold | Conditional Random Field | $O(n^3)$ | slowest |
| MXfold | structured SVM | $O(n^3)$ | medium |
| Our work | structured SVM | LinearFold | fastest |

## 2 Methods

### 2.1 Structured SVM Training with Linear-Time Inference

We use structured SVM (sSVM) for training and LinearFold for linear-time inference. This makes the training process faster than both CONTRAfold and MXfold, especially when the training dataset contains long sequences.

Formally, we define 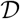 as the training dataset, and (*x*, *y*) as RNA sequence and its structure in 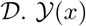 is the set of all possible structures for the given sequence *x*. *y*′ is a predicted structure, i.e., 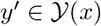. sSVM training is to learn a feature weight vector **w**, which scores the native structure *y* higher than *y*′ by a margin of *Δ*(*y*, *y*′):

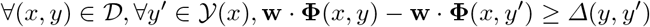

where **Φ** is the feature function, mapping (*x*, *y*) (or (*x*, *y*′)) to its feature vector. *Δ*(*y*, *y*′) is the decomposable augmented loss to measure the difference between *y* and *y*′. We define *Δ*(*y*, *y*′) as:

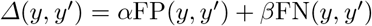

where FP(*y*, *y*′) is the number of False Positive pairs and FN(*y*, *y*′) is the number of False Negative pairs. *α* and *β* are hyper-parameters that balance PPV and sensitivity.

**Algorithm 1.**
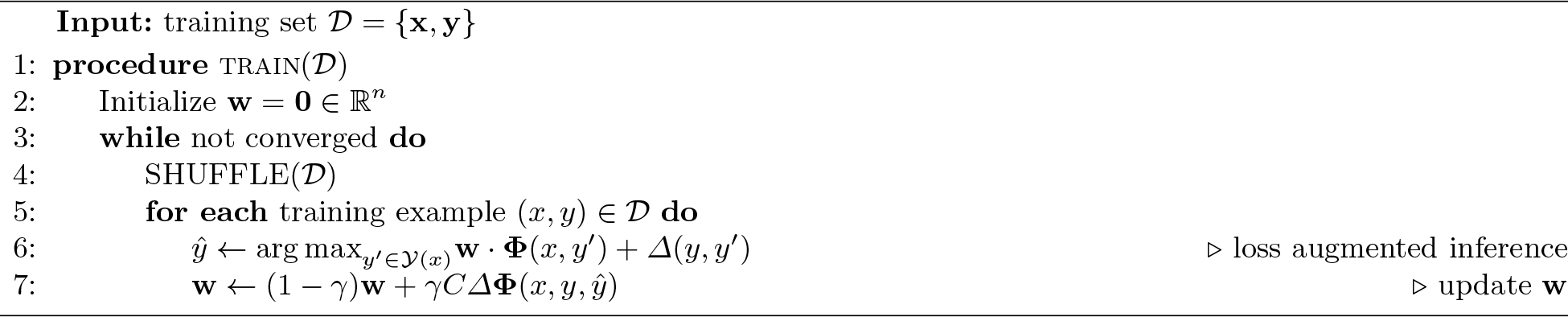
structured SVM training with SGD

For simplicity, we define *Δ***Φ**(*x*, *y*, *y*′) as the feature vector difference of *y* and *y*′, i.e., *Δ***Φ**(*x*, *y*, *y*′) : = **Φ**(*x*, *y*) − **Φ**(*x*, *y*′). A triple (*x*, *y*, *y*′) is called to be a *violation* with respect to **w** if:

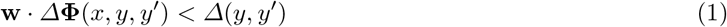

Among all the violations, we define the most violated structure 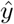:

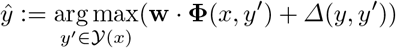

Previous work by MXfold gets 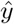 with loss-augmented decoding through exact search, but since the exact search-based inference requires *O*(*n*^3^) runtime, the inference is slow for long sequences, leading to a costly training process. Alternatively, we use LinearFold as a linear-time inference engine to get *y*\*, an approximation of 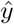:

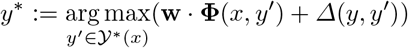

where 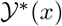 is the search space of LinearFold and 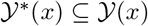.

### 2.2 Convergence Analysis for sSVM with Inexact Search

Though the training process can be accelerated with linear-time inexact search-based inference, the search error may lead to invalid updates and result in non-convergence. Next, we will show why sSVM’s convergence property does not hold when applying inexact search.

First, we review the condition of exact search and analyze its convergence. Algorithm 1 presents the pseudocode of naive structured SVM training via Stochastic Gradient Descent (SGD), using exact search for loss augmented inference (line 6). We can prove that for a separable dataset sSVM training will make finite number of updates (before convergence) when doing exact search for inference.

#### Definition 1.

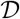 *is structured SVM separable with a margin of* Δ(*y*, *y*′)*, if there exists an oracle feature weight vector* **u** *with* ||**u**|| = 1, *s.t. it can correctly classify all examples in* 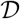 *with a gap of at least* Δ(*y*, *y*′). *maximal margin* 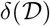 *is the maximal such margin over all unit oracle vectors*:

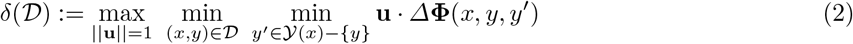

It is clear that 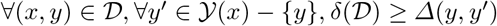.

#### Definition 2.

*The diameter R* 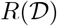 *of* 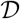 *is*:

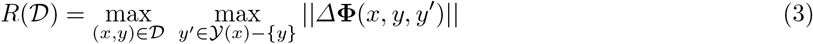

#### Theorem 1.

*For a structured SVM separable dataset* 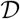 *with margin* Δ(*y*, *y*′), *the stochastic gradient descent (SGD) algorithm (see Algorithm 1) will make finite number of updates (before convergence)*.

*Proof*. Training structured SVM via SGD is to minimize:

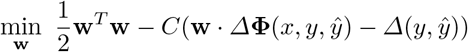

The subgradient of the objective function is:

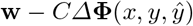

Denote **w**^**k**^ to be the weight vector at step *k*, and *γ* to be the learning rate. At each step, **w** is updated as:

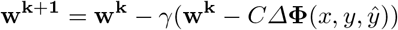

We bound ||**w**^**k+1**^|| in two directions. For detailed proof please refer to supporting information.

1. Upper bound:

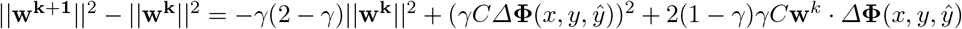 Since the update is because of a violation, by Equation 1 we have (*C* > 0):

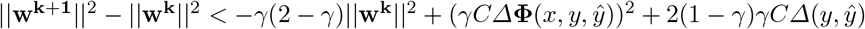 If we choose 0 < *γ* < 1, by induction and Definition 1, 2, we can get the upper bound of:

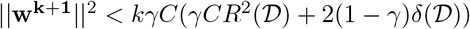
2. Lower bound:

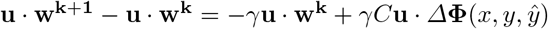

By Definition 1 and upper bound of ||**w**^**k**^||, we get the lower bound as:

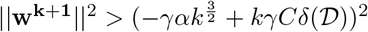

where 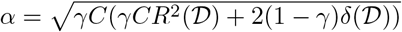.
3. Combine upper and lower bound together, we have:

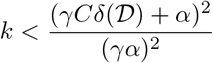

The key step in upper bound part is that the update from **w**^**k**^ to **w**^**k**+**1**^ is because of a violation. When the inference uses exact search, the most violated structure 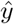 is guaranteed to score higher than ground truth structure *y*. However, when the search is inexact, for example beam search in LinearFold, it is possible that ground truth structure is not in the search space (i.e., 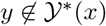). Thus, even if the predicted structure *y*\* is different from *y*, it can still score lower, and the update is therefore invalid. When we do an invalid update,we shift **w** towards a wrong direction and the convergence may not hold.

**Algorithm 2.**
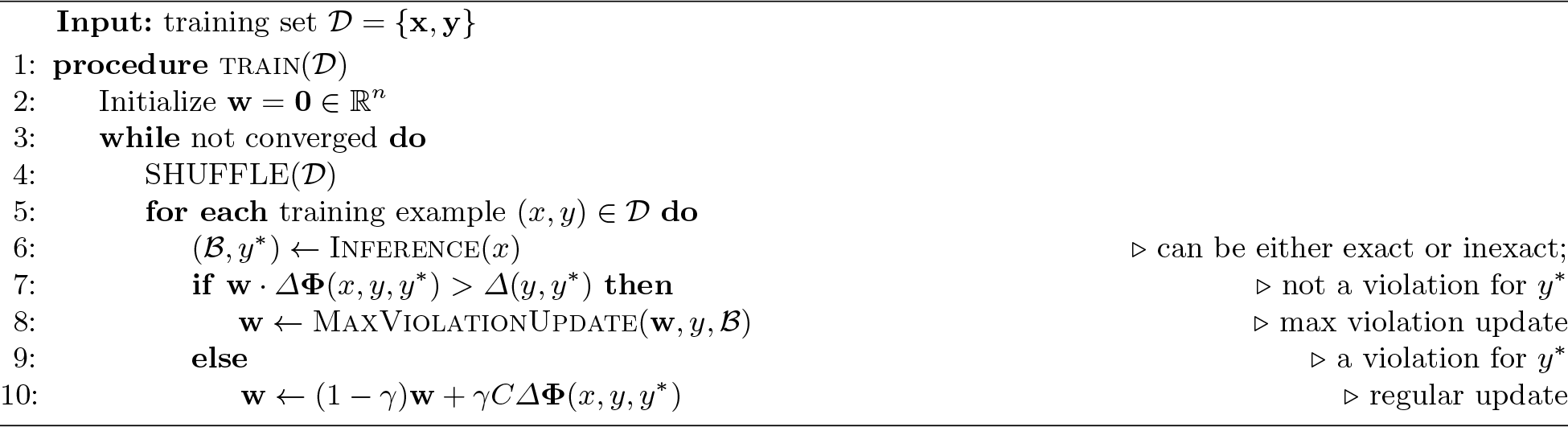
structured SVM with Inexact Search Inference and Max Violation Update

### 2.3 Max Violation Update Strategy

To remedy the non-convergence introduced by inexact search inference, We now propose the new framework of sSVM training, which uses max violation update strategy and can converge with inexact search.

The target of sSVM training is to learn a feature weight vector **w**, which scores ground truth structure *y* higher than any other structure, i.e., no violation for all examples. So, the main observation is each update is due to a violation. For exact search inference, if the predicted structure 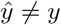, it must be a violation and 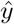 must be the most violated structure. However, for inexact search inference, even if the predicted structure *y*\* ≠ *y*, it is not necessary a violation since it is possible for *y* to have a higher score than *y*\*.

With this observation, it is clear that the key solution is to make sure each update is for a violation, no matter if the search is exact. We introduce a max violation update strategy, which is a successful strategy in structured perceptron training. Basically, this strategy uses the most violated prefix of the ground truth structure and predicted structure for update. More formally, denote *y*_mv_ and 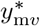 as the max violation prefix of *y* and *y*\*, and define them as:

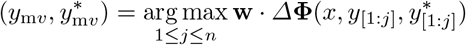

where *n* is the sequence length and *j* is each position in the sequence. *y*_[1:*j*]_ is the prefix of *y* up to *j*. Note that 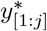 denotes the predicted substructure with the highest score up to *j* in the searching space, and 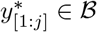, where 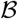 is the set of all such predicted substructures for every position *j*.

Algorithm 2 presents the pseudocode of sSVM training with max violation updated strategy. Now in line 6 it does not require an exact search (i.e., arg max) for inference, for example, we can use LinearFold as the inference engine. Then we check if *y*\* results in a violation. If not, we do max violation update. Otherwise, we do regular update as in Algorithm 1.

## 3 Results

### 3.1 Training Time

First, we test the training time of our system, and compare with two machine learning-based systems, CONTRAfold and MXfold. We use ArchiveII dataset [16,18] ^1^, which contains 3,857 RNA sequences from 10 different RNA families. The average and max length of the sequences are 208.6 and 2,968. The 10 families (listed from the shortest to the longest) are tRNA, 5S rRNA, SRP, RNaseP, tmRNA, Group I Intron, Group II Intron, telomerase RNA, 16S rRNA and 23S rRNA.

Figure 1A compares per iteration runtime result among all three systems. We can see that CONTRAfold is the slowest and takes about 290 minutes, while MXfold is faster and takes about 15.37 minutes. Our system is the fastest and takes only 7.06 minutes, which is 41× faster than CONTRAfold training and 2.2× faster than MXfold training.

**Fig. 1:**
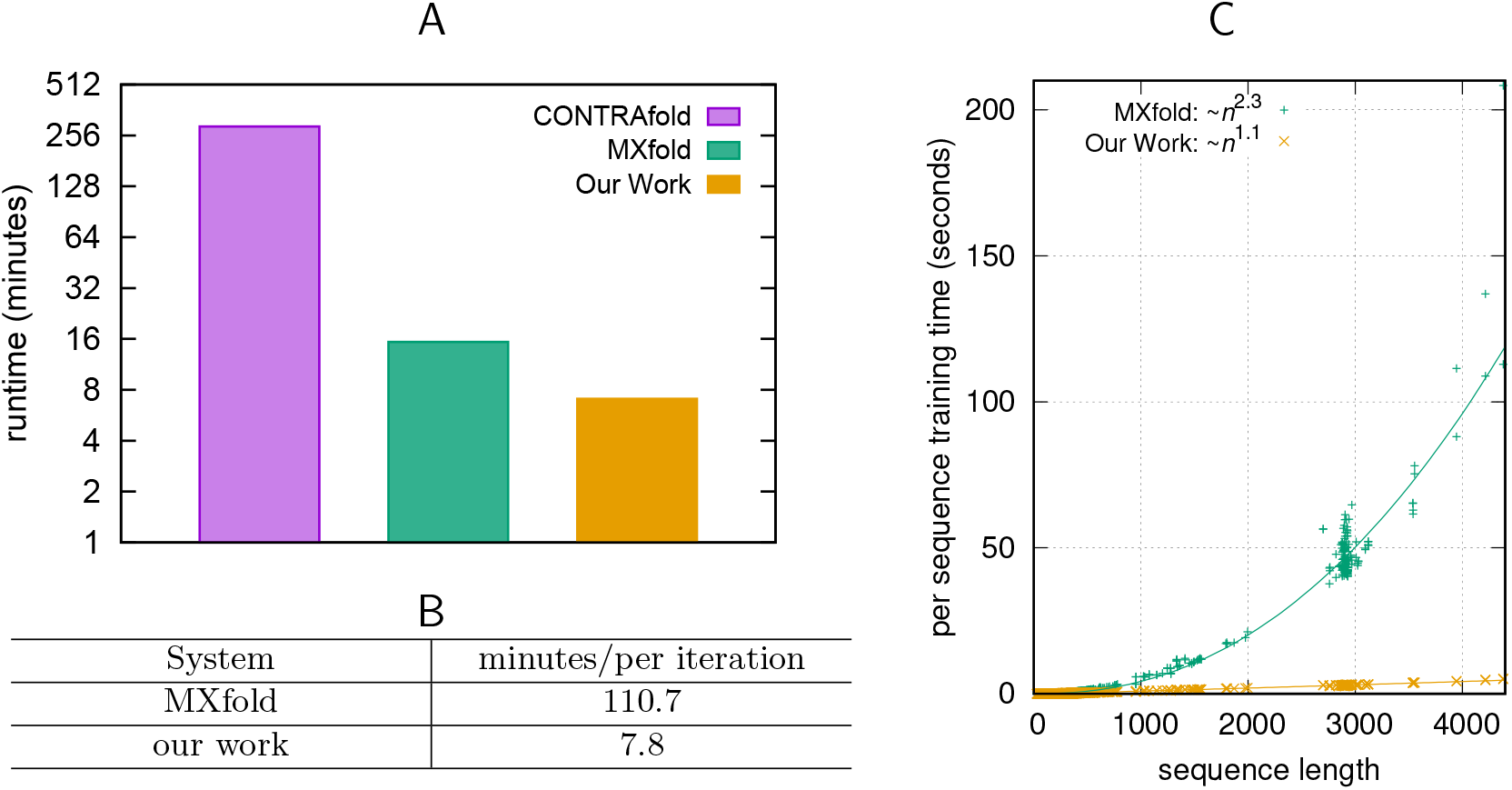
Training runtime comparison between our system and the baselines. **A**: per iteration training runtime comparison between CONTRAfold, MXfold and our system on ArchiveII dataset. Note that the *y*-axis is in log scale. **B**: per iteration training runtime comparison between MXfold and our system on 23S rRNA dataset from bpRNA. **C**: per sequence training runtime comparison between MXfold and our system on these two datasets.

The speedup made by MXfold is mainly because sSVM training algorithm is faster than CRF used by CONTRAfold. Since MXfold still uses cubic algorithm for inference, we infer that if the dataset contains more long sequences, MXfold training will be much slower. To verify this, we collect long sequences in 23S rRNA family from bpRNA dataset [5] ^2^, and get a dataset containing 163 sequences, with average length 2713.5*nt*. Figure 1B shows the per iteration runtime comparison between MXfold and our system on this longer dataset. Since CONTRAfold is too slow, we do not include it. We can see that our system is about 14× faster, which comfirms that our system is much faster for training on longer sequences.

In addition, we test the runtime for each sequence to verify the linearity of our system. Figure 1C presents such comparison between MXfold and our system. We can see that our system scales almost linearly with the sequence length, while MXfold has super-quadratic runtime. The minor deviations from the theoretical runtimes are due to the majority of short sequences in ArchiveII dataset. For the longest sequence with length about 4381*nt*, MXfold takes about 208 seconds while our work only takes about 5 seconds, which is about 41× faster. We also notice that the runtime deviations of sequences with similar length for MXfold are much larger than our system, which suggests that our system can finish training within an accurate estimated time.

### 3.2 Accuracy

Next, we compare accuracy with both thermodynamics-based system and machine learning-based systems. We use cross-validation (leave-one-out) training on ArchiveII dataset, i.e., train on 9 families and test on the other family, to verify the learned models have strong generalization ability. We report Positive Predictive Value (PPV, the fraction of predicted pairs in the known structure) and sensitivity (the fraction of known pairs predicted) for each family, and the overall PPV, sensitivity and F1-score 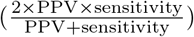 which are averaged over families. Following previous slipping method, we allow base pair to slip by one nucleotide [18]. MXfold extends CONTRAfold features, and we also add these extra features into LinearFold when using it as the inference engine.

Figure 2 shows the per family and overall accuracies. We use 5 systems as baselines, the off-the-shelf versions of CONTRAfold, MXfold, LinearFold-C (LinearFold with machine learning-based model) and LinearFold-V (LinearFold with thermodynamics-based model), as well as retrained MXfold. CONTRAfold is too slow for retraining, so we do not include retrained CONTRAfold as a baseline. We do cross-validation training for our system and MXfold for 100 epochs.

**Fig. 2:**
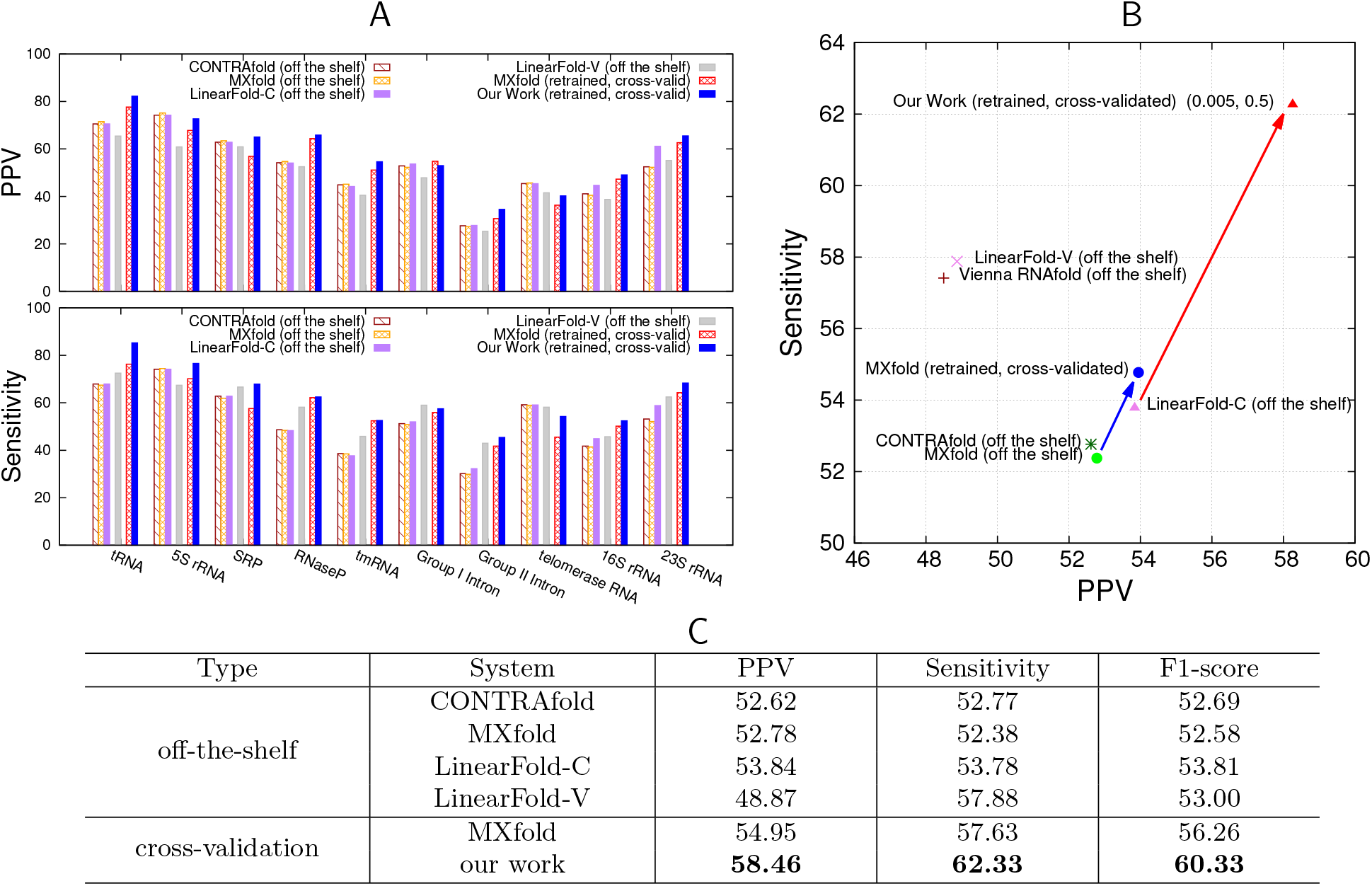
Accuracy comparison between different systems. **A**: per family PPV and sensitivity comparison. **B**: overall PPV-sensitivity-tradeoff comparison. (0.005, 0.5) are the values of hyper-parameters (*α, β*). **C**: table for overall PPV, sensitivity and F1-score, averaging over all families.

Figure 2A shows that our system has the highest PPV on all families except 5S rRNA, Group I Intron and telomerase RNA, as well as the highest sensitivity on all families except Group I Intron and telomerase RNA. Note that off-the-shelf machine learning-based systems may produce better results as they may have corresponding families in their training sets. Our system outperforms retrained MXfold on both PPV and sensitivity for all families, except for PPV of Group I Intron. The results verify that our linear-time training system can learn better models, and also shows that our system has a good generalization ability of learning models which can be used across families.

We demonstrate PPV-sensitivity-tradeoff in figure 2B, and show our work’s PPV-sensitivity with selected hyper-parameters (*α* = 0.005, *β* = 0.5). We can see that our system achieves the highest overall PPV and sensitivity among all systems, including machine learning-based systems and thermodynamics-base systems, by big margins. Figure 2C gives the exact number of overall PPV, Sensitivity and F1-Score in different systems. Our system improves off-the-shelf LinearFold-C by +4.62% for PPV, +8.55% for sensitivity and +6.51% for F1-score, and outperforms retrained MXfold by +3.51%, +4.70%, +4.07% on PPV, sensitivity and F1-Score, separately. Compared with some other off-the-shelf systems, the improvement is even bigger. For example, Our system outperforms the off-the-shelf CONTRAfold by +5.84%, +9.56%, +7.64% on PPV, sensitivity and F1-Score, separately.

We borrow MXfold feature template for all experiments in figure 2, but we notice that even with the basic feature template in CONTRAfold (about 336 non-zero features), our system achieves a similar accuracy as using MXfold feature template (around 4,000 non-zero features). Table 2 shows the accuracy comparison for systems with different feature templates. We can see that PPV and sensitivity changes of our system with different feature template are very small, only 0.11% and 0.01%, separately. In addition, even with a much smaller feature template, our system is better than MXfold. On the other hand, MXfold integrates thermodynamic model in its system, which helps for generalization and makes it a mixture of learning-based and thermodynamics-base system. If thermodynamic model is disabled and train for a pure learning model for MXfold, the PPV and sensitivity drop −0.53% and −2.81%, separately. Compared to this pure learning-based MXfold, our system’s accuracy improvement is even more salient.

**Table 2:** Accuracy comparison of systems with different feature templates.

| Features | System | PPV | Sensitivity | F1-score |
| --- | --- | --- | --- | --- |
| basic | CONTRAFold (off-the-shelf) | 52.62 | 52.77 | 52.69 |
|  | our work (retrained) | **58.35** | **62.32** | **60.27** |
| rich | MXfold (retrained, w/o thermodynamic model) | 54.42 | 54.82 | 54.62 |
|  | MXfold (retrained, w/ thermodynamic model) | 54.95 | 57.63 | 56.26 |
|  | our work (retrained) | **58.46** | **62.33** | **60.33** |

### 3.3 Update Strategy Impact

In section 2.3 we introduce max violation update strategy as a fix for invalid update in sSVM training with inexact search inference. Here we investigate the impact of different update strategies, i.e., regular update and max violation update, on accuracy.

Table 3 shows the accuracy changes between regular and max violation update strategies on ArchiveII cross validation training. Since the search quality is highly related to beam size, we show the comparison between different beam sizes too. We can see that with small beam sizes *b* = 1 and *b* = 5, the search quality is bad, thus max violation update strategy leads to a better accuracy. With beam size increasing to *b* = 50 or even *b* = 100, the search quality for most short sequences is good (i.e., doing exact search for families such as tRNA and 5S rRNA), and invalid update becomes fewer, resulting in closer accuracies between two different update strategies. This confirms invalid updates are the key for sSVM training with inexact search.

**Table 3:**
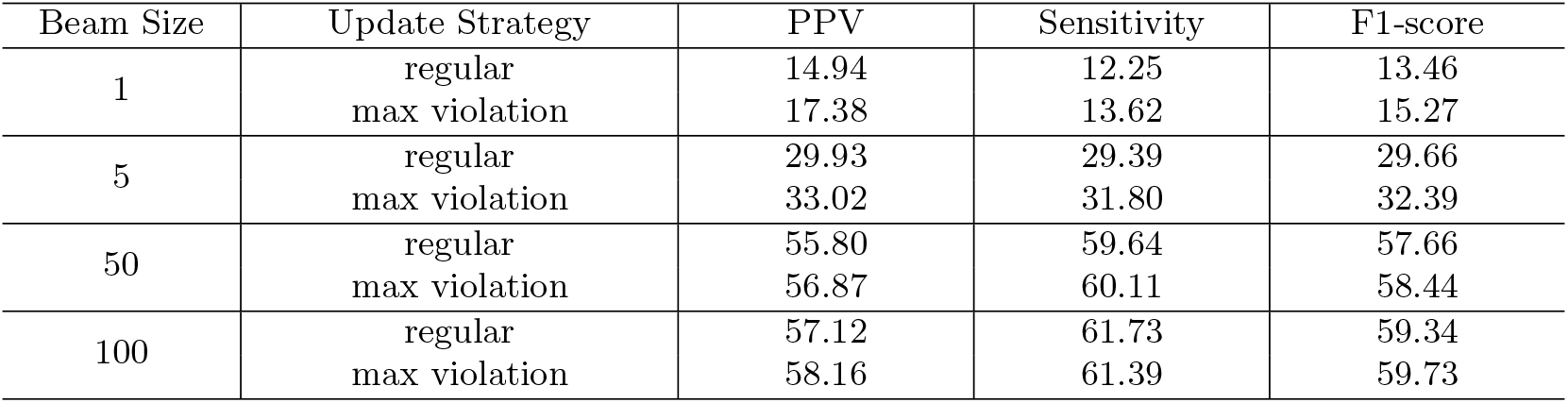
Comparison between regular and max violation update strategy for different beam sizes on ArchiveII cross validation training (run for 10 epochs).

Since ArchiveII dataset is very diverse, which contains both very long (about 2,900 *nt*) and very short (about 50 *nt*) sequences, it is hard to choose a balanced beam size for training, i.e., a small beam size is suitable for short families but bad for long families, and a large beam size makes the search for most sequences become exact search. However, on some other less diverse dataset, for example S-Processed dataset (a data set used by CONTRAfold v2.02 training, originally from S-full dataset [2] ^3^), the impact is more salient.

Figure 3 shows the impact of different update strategy on S-Processed dataset training. Figure 3A shows that for beam size *b* = 5, max violation update strategy leads to a better accuracy. Compared to regular update, max violation update helps to gain about +5% increase on F1-score. In addition, the curve of regular update drops after 500 seconds, but max violation update curve keeps increasing in all the training process. Figure 3B visualizes an example from S-Processed test set. This example’s length is 106*nt*, and is part of a 16S rRNA sequence. We show three structures of this example, ground truth structure, regular update structure and max violation update structure. We can see that prediction structure with max violation update strategy is almost the same as the ground truth structure, and its PPV and sensitivity are 96.97% and 94.12%, separately. Not as good as max violation update structure, prediction structure with regular update strategy is very different from the ground truth structure, which incorrectly predicts all of the base pairs.

**Fig. 3:**
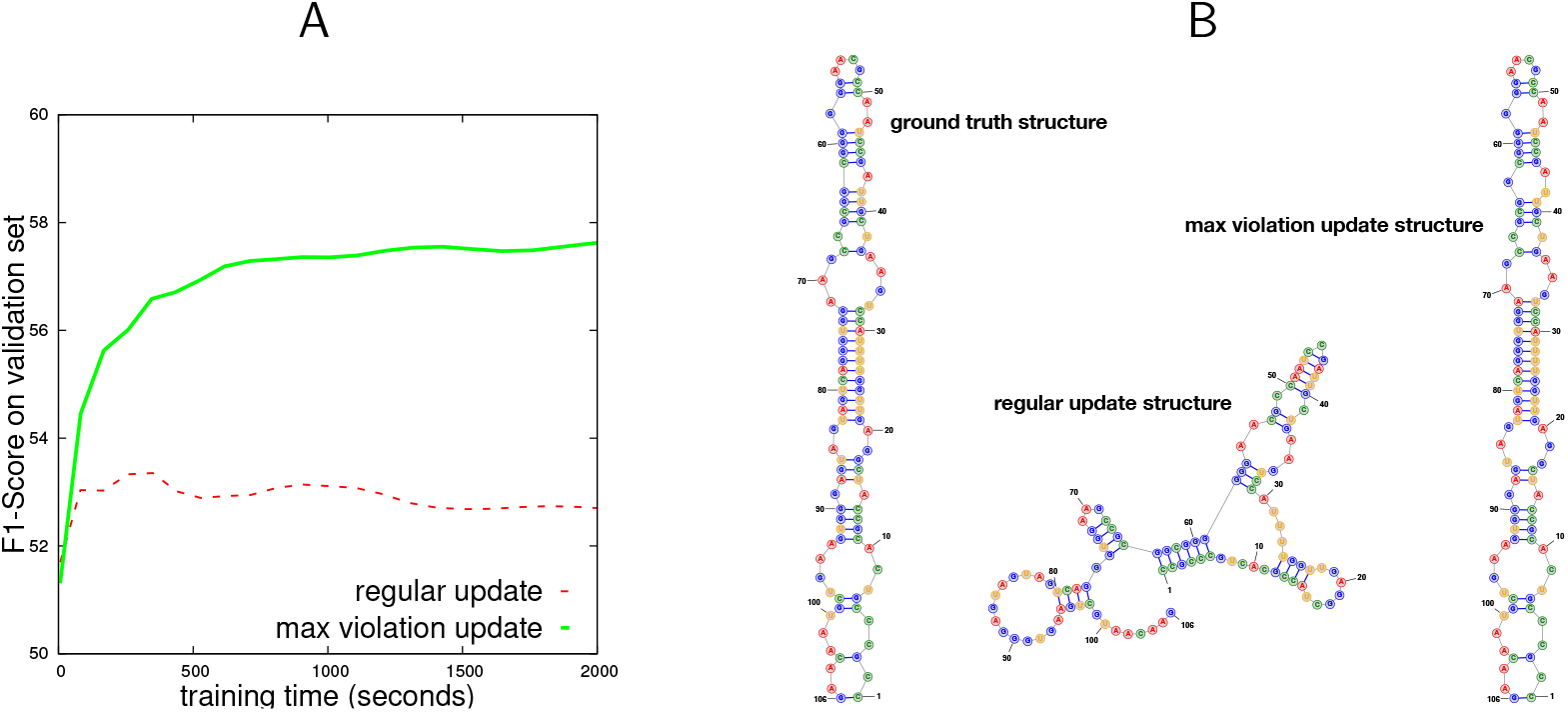
Impact of different update strategy on S-Processed dataset training. **A**: F1-score curves of training processes with two different update strategies for beam size 5. **B**: An example showing that max violation update strategy gives a more accurate structure prediction. RNA secondary structures are drawn with StructureEditor.

## 4 Discussion

Machine learning-based methods for RNA secondary structure prediction can learn feature weights from known structures, and address the inaccurate measurement of thermodynamic parameters due to experimental limitation. However, training with big datasets containing diverse sequences, which is beneficial for learning-based methods, is either impossible or very expensive due to the slowness of current learning systems. To address this issue, we propose a new machine learning-based system, which can learn to fold in linear time. Our system uses the successful linear-time prediction engine LinearFold as inference, and uses structured SVM, a much faster machine learning algorithm than CRF used by CONTRAfold, for training procedure. Furthermore, we utilize a max violation update strategy to address the non-convergence issue introduced by using LinearFold as inexact search inference, and generalize the naive sSVM algorithm to a max violation update version of sSVM.

We confirm that:

1. The training speed of our system is 41× faster than CONTRAfold for one epoch even on a diverse dataset (average length 208.6 *nt*). On a dataset with more long sequences (average length 2713.5 *nt*), our system is 14× faster than MXfold. The training time for each sequence increases linearly with sequence length. See figure 1.
2. The accuracy of our system outperforms both thermodynamics-based systems and machine learningbased systems, and achieves the highest accuracy in most families. Our system has a strong generalization ability, which can learn good models from cross-validation training. See figure 2.
3. Max violation update strategy leads to accuracy improvement. See figure 3.

Our system can be extended to deep learning-based RNA folding system. Now our model still uses manually designed features, but it can also be built on some well-designed neural networks, which can automatically extract implicit features. With these features, it is possible to get a more accurate RNA folding prediction system.

## Footnotes

1 http://rna.urmc.rochester.edu/pub/archiveII.tar.gz

2 http://bprna.cgrb.oregonstate.edu/

3 http://www.rnasoft.ca/CG/

