## Supporting Information for "Learning to Fold RNAs in Linear Time"

### Supporting Information
### Learning to Fold RNAs in Linear Time

**F A Rezaur Rahman Chowdhury, He Zhang and Liang Huang**

#### 1 sSVM Convergence Proof

*Proof.* Training structured SVM via SGD is to minimize:

$$\min_{\mathbf{w}} \frac{1}{2}\mathbf{w}^T\mathbf{w} - C(\mathbf{w} \cdot \Delta\mathbf{\Phi}(x, y, \hat{y}) - \Delta(y, \hat{y}))$$

The subgradient of the objective function is:

$$\mathbf{w} - C\Delta\mathbf{\Phi}(x, y, \hat{y})$$

Denote $\mathbf{w^k}$ to be the weight vector at step $k$, and $\gamma$ to be the learning rate. At each step, $\mathbf{w}$ is updated as:

$$\mathbf{w^{k+1}} = \mathbf{w^k} - \gamma(\mathbf{w^k} - C\Delta\mathbf{\Phi}(x, y, \hat{y}))$$

We will bound $||\mathbf{w^{k+1}}||$ from two directions.

1. First let's analyze the upper bound.

$$||\mathbf{w^{k+1}}||^2 = ||\mathbf{w^k} - \gamma(\mathbf{w^k} - C\Delta\mathbf{\Phi}(x, y, \hat{y}))||^2$$
$$\Longrightarrow$$
$$||\mathbf{w^{k+1}}||^2 - ||\mathbf{w^k}||^2 = -\gamma(2-\gamma)||\mathbf{w^k}||^2 + (\gamma C\Delta\mathbf{\Phi}(x, y, \hat{y}))^2 + 2(1-\gamma)\gamma C\mathbf{w}^k \cdot \Delta\mathbf{\Phi}(x, y, \hat{y})$$

Since the update is because of a violation, from Equation 1 we have ($C > 0$):

$$||\mathbf{w^{k+1}}||^2 - ||\mathbf{w^k}||^2 < -\gamma(2-\gamma)||\mathbf{w^k}||^2 + (\gamma C\Delta\mathbf{\Phi}(x, y, \hat{y}))^2 + 2(1-\gamma)\gamma C\Delta(y, \hat{y})$$

Then if we set $0 < \gamma < 1$, and combine with Definition 1, we have:

$$\begin{aligned} ||\mathbf{w^{k+1}}||^2 - ||\mathbf{w^k}||^2 &< (\gamma C\Delta\mathbf{\Phi}(x, y, \hat{y}))^2 + 2(1-\gamma)\gamma C\Delta(y, \hat{y}) \\ &\leq (\gamma C\Delta\mathbf{\Phi}(x, y, \hat{y}))^2 + 2(1-\gamma)\gamma C\delta(\mathcal{D}) \end{aligned}$$

By Definition 2 and induction, we have the upper bound for $||\mathbf{w^{k+1}}||$:

$$\begin{aligned} ||\mathbf{w^{k+1}}||^2 &< k(\gamma C\Delta\mathbf{\Phi}(x, y, \hat{y}))^2 + 2k(1-\gamma)\gamma C\delta(\mathcal{D}) \\ &\leq k\gamma C(\gamma CR^2(\mathcal{D}) + 2(1-\gamma)\delta(\mathcal{D})) \end{aligned}$$

2. Then let's analyze the lower bound.

$$\mathbf{u} \cdot \mathbf{w^{k+1}} = (1-\gamma)\mathbf{u} \cdot \mathbf{w^k} + \gamma C\mathbf{u} \cdot \Delta\mathbf{\Phi}(x, y, \hat{y})$$
$$\Longrightarrow$$
$$\mathbf{u} \cdot \mathbf{w^{k+1}} - \mathbf{u} \cdot \mathbf{w^k} = -\gamma\mathbf{u} \cdot \mathbf{w^k} + \gamma C\mathbf{u} \cdot \Delta\mathbf{\Phi}(x, y, \hat{y})$$

By Definition 1, we have:

$$\mathbf{u} \cdot \mathbf{w^{k+1}} - \mathbf{u} \cdot \mathbf{w^k} \geq -\gamma \mathbf{u} \cdot \mathbf{w^k} + \gamma C \delta(\mathcal{D})$$

$\mathbf{u} \cdot \mathbf{w^k}$ have upper bound as:

$$\mathbf{u} \cdot \mathbf{w^k} \leq ||\mathbf{u}||||\mathbf{w^k}||$$

Since we have upper bound for $||\mathbf{w^k}||$, let $\alpha = \sqrt{\gamma C(\gamma C R^2(\mathcal{D}) + 2(1-\gamma)\delta(\mathcal{D}))}$ and $\alpha > 0$, so we have:

$$\begin{aligned}
\mathbf{u} \cdot \mathbf{w^k} &\leq ||\mathbf{u}||||\mathbf{w^k}|| \\
&\leq \alpha(k-1)^{\frac{1}{2}} \\
&< \alpha k^{\frac{1}{2}}
\end{aligned}$$

So,

$$\begin{aligned}
\mathbf{u} \cdot \mathbf{w^{k+1}} - \mathbf{u} \cdot \mathbf{w^k} &> -\gamma \alpha k^{\frac{1}{2}} + \gamma C \delta(\mathcal{D}) \\
\implies \mathbf{u} \cdot \mathbf{w^{k+1}} &> -\gamma \alpha \sum_i^k i^{\frac{1}{2}} + k \gamma C \delta(\mathcal{D}) \\
&> -\gamma \alpha k^{\frac{3}{2}} + k \gamma C \delta(\mathcal{D}) \\
||\mathbf{u}||^2 ||\mathbf{w^{k+1}}||^2 &> (-\gamma \alpha k^{\frac{3}{2}} + k \gamma C \delta(\mathcal{D}))^2 \\
||\mathbf{w^{k+1}}||^2 &> (-\gamma \alpha k^{\frac{3}{2}} + k \gamma C \delta(\mathcal{D}))^2
\end{aligned}$$

3. Combine with the upper bound and lower bound, we have:

$$(-\gamma \alpha k^{\frac{3}{2}} + k \gamma C \delta(\mathcal{D}))^2 < \alpha^2 k$$

Since k is a positive integer, we have:

$$\begin{aligned}
(-\gamma \alpha k^{\frac{3}{2}} + k \gamma C \delta(\mathcal{D}))^2 &< \alpha^2 k^2 \\
-\alpha k < -\gamma \alpha k^{\frac{3}{2}} + k \gamma C \delta(\mathcal{D}) &< \alpha k \\
\implies \gamma \alpha k^{\frac{1}{2}} &< \gamma C \delta(\mathcal{D}) + \alpha \\
\implies k &< \frac{(\gamma C \delta(\mathcal{D}) + \alpha)^2}{(\gamma \alpha)^2} \\
k &< \frac{(\gamma C \delta(\mathcal{D}) + \alpha)^2}{(\gamma \alpha)^2}
\end{aligned}$$

where $\alpha = \sqrt{\gamma C(\gamma C R^2(\mathcal{D}) + 2(1-\gamma)\delta(\mathcal{D}))}$.
